# Lineage tracing reveals *atoh7* positive and negative retinal ganglion cell populations in the zebrafish retina

**DOI:** 10.64898/2026.03.19.712911

**Authors:** Darby M. Bennett, Robert I. Newland, Matthew B. Veldman, Joel B. Miesfeld

## Abstract

**Purpose:** *Atoh7* is a transiently expressed developmental transcription factor that gives rise to the seven major retinal cell types. Despite this broad lineage, *Atoh7* is only required for retinal ganglion cell (RGC) genesis and survival, even though a significant portion of RGCs are *Atoh7* negative based on lineage tracing in mice, suggesting a cell nonautonomous role for *Atoh7* in the genesis and survival of all RGCs. Atoh7 function is conserved in zebrafish, yet the full retinal lineage, including the RGC population, has remained unidentified. Therefore, we sought to determine the *atoh7* retinal lineage in wild type and *atoh7* mutant zebrafish retinas.

**Methods:** We generated *atoh7*:iCre transgenic zebrafish and in combination with the established *ubi*:Switch lineage trace permanently labeled cells that represent the *atoh7* lineage. A combination of *in vivo* live imaging and immunohistochemical techniques were used to validate *atoh7*:iCre transgene expression and the *atoh7* lineage in embryonic, larval, and adult retinas as well as the adult brain.

**Results:** The *atoh7*:iCre;*ubi*:Switch transgene combination successfully recapitulated the onset of endogenous *atoh7* expression and transgene fluorophores persisted into adulthood labeling the *atoh7* lineage. Most notably, we determined 79% of total RGCs in the wild type retina come from *atoh7+* progenitor cells, a greater number than reported in the mouse retina. In *atoh7* mutant retinas, we confirmed a complete loss of RGCs and observed a statistically significant increase in the proportion of *atoh7*+/Pax6+ amacrine cells, as well as an increase in the total number of Prox1+ bipolar cells. Interestingly, we discovered *atoh7*+ cells located outside the eye in other areas of the central nervous system.

**Conclusions:** These data demonstrate the presence of *atoh7* positive and negative retinal cell types in the zebrafish retina, including RGCs, highlighting the potential to study survival mechanisms of *atoh7* negative RGCs and fate switch paradigms using zebrafish retinal development models.

## Introduction

The vertebrate neural retina is composed of seven major cell types, including amacrine, bipolar, horizontal, retinal ganglion cells (RGCs), Müller glia (MG), and rod and cone photoreceptors, which arise from a common pool of multipotent naïve retinal progenitor cells (RPCs)^1–4^. Retinal neurogenesis begins with naïve RPC expression of a combination of pro-neurogenic transcription factors, transitioning RPCs into each of the post mitotic retinal cell types. One of these pro-neurogenic factors is *Atoh7* (*drosophila* atonal homolog 7), a basic-helix-loop-helix transcription factor that begins expressing prior to the onset of retinal neurogenesis, with its lineage representing all seven major retinal cell types^5^. Despite its broad lineage, *Atoh7* is only essential for the genesis and survival of RGCs, the first-born retinal cell type. In all species studied, loss of *Atoh7* results in a ≥95% loss of all RGCs, while the other retinal cell types are either modestly or completely unaffected^5–10^. Interestingly, based on lineage tracing only 55% of total RGCs in the adult mouse retina come from the *Atoh7+* lineage despite the severe loss of RGCs in *Atoh7* mutant retinas^5^. These data suggest the *Atoh7+* RPC or RGC population may be required for the survival of all RGCs, although the exact mechanism remains unknown. Zebrafish are an ideal model to study this complex interaction, but it has not been established if they contain an *atoh7* negative RGC lineage.

*Atoh7* is transiently expressed during development, with its mRNA and protein being undetectable in fully differentiated retinal neurons, thus making it difficult to determine the full *atoh7* lineage^5,11–14^. In fish, *atoh7* (previously *ath5*) expression begins at ∼25 hours post-fertilization (hpf) in the ventro-nasal retina. It spreads dorso-nasally and ventro-temporally, preceding the wave of retinal neurogenesis, and completes this wave by 48 hpf^12,14^. By 72 hpf, *atoh7* mRNA is restricted to the ciliary margin^14^. The first tracking of the zebrafish *atoh7* retinal lineage was done using an *atoh7* promoter driven transgene, *ath5*:GFP^12,15,16^. The stability of GFP allowed for successful tracking of the *atoh7* lineage, revealing a faithful expression pattern during early neurogenesis and post mitotically in RGCs, photoreceptors, amacrine, and horizontal cells at 4 days post-fertilization (dpf)^12,15,17^. Despite the ability to track the *atoh7* lineage at these early timepoints, the *ath5*:GFP transgene is limited in its ability to determine the *atoh7* lineage of the entire neural retina at late stages when all cell types are born.

To determine the zebrafish *atoh7* RGC and full retinal lineage in the mature retina, we created a transgenic line utilizing the zebrafish *atoh7* promoter to drive expression of Cre recombinase, *atoh7*:iCre. *atoh7* lineage+ cells were permanently labeled by combining the *atoh7*:iCre transgene with the *ubi*:Swtich (LoxP-eGFP-LoxP-mCherry) transgene^18,19^. mCherry expression matched the hallmark spatiotemporal expression pattern of endogenous *atoh7* and was detected into adulthood^14,17,18^. We determined there are both *atoh7* lineage positive and negative populations of RGCs, and all other major retinal cell types are represented in the *atoh7* lineage, albeit in differing percentages compared to mice. We further characterized the *atoh7* lineage in *atoh7* (*lakritz)* mutants^6^, discovering that the proportion of *atoh7* lineage+ populations remain the same in all cell types, except RGCs and amacrine cells. Lastly, *atoh7* is expressed in cell populations within the central nervous system outside of the neural retina. The data presented indicate that our lineage tracing transgenes create an innovative way to study the possible genesis and survival mechanisms between the positive and negative *atoh7* lineage cell populations of RGCs, other major retinal cell types, and a variety of cells in the CNS.

## Methods

### Zebrafish Husbandry

All adult zebrafish were housed in a controlled environment with a water temperature of 28°C and 14h light/10h dark cycle. Embryos were raised at 28°C in 1x InstantOcean (IO) Sea salt (0.06 g/L) upon collection and transferred at 24 hpf to 1xIO/1X PTU (0.003% 1-phenyl-2-thiourea, Acros Organics/Thermo Fisher Chemicals) to prevent pigmentation for transgenic screening, then returned to 1x IO until 5 dpf. For *in* vivo imaging, zebrafish larvae were anesthetized with 1x Tricaine (0.17 g/L, Tokyo Chemical Industry Co., Ltd.) diluted in 1x IO/1X PTU. Animals were euthanized with cold 1x Tricaine diluted in 1x IO/1X PTU. All animal work was performed in accordance with the Institutional Animal Care and Use Committee of the Medical College of Wisconsin, ARVO Statement for the Use of Animals in Ophthalmic and Vision Research, and National Institutes of Health Guide for the Care and Use of Laboratory Animals.

### Plasmid and Transgenic Line Generation

The *atoh7:*iCre*;cmlc:*eGFP (*atoh7*:iCre) plasmid was generated using the Gateway and the Tol2 kit to combine the previously published *atoh7* (*ath5*) zebrafish promoter^12,19^ and improved Cre recombinase (iCre)^21^ in the pDEST vector backbone including *cmlc*:eGFP for screening^21^. *Tol2 transposase* mRNA (50pg) was co-injected with the completed *atoh7*:iCre plasmid (50pg) in 1-4 cell stage zebrafish embryos^22^. All experiments were conducted on the second generation (F2) to verify single insert inheritance and expression pattern consistency. The *atoh7*:iCre line is designated as *mw800* in The Zebrafish Information Network database (zfin.org).

### Immunohistochemistry

Larval zebrafish and adult heads were fixed in 4% paraformaldehyde (PFA)/1X phosphate buffered saline (PBS) overnight at 4°C, washed 3 times in 1X PBS, and processed through a sucrose gradient (5% or 15% sucrose/1X PBS/0.2% Na azide) prior to cryopreservation in O.C.T. (optimal cutting temperature medium, Thermo Scientific). Serial sections (10μm) were collected and rehydrated with 1X PBS, treated with a MeOH series (33%, 66%, 100%) to reduce endogenous eGFP and mCherry fluorescence from the *ubi*:switch transgene followed by 3 washes in 1X PBS. Standard immunohistochemistry with 4% (w/v) nonfat dry milk and 0.1% (w/v) Tween-20 in Tris-buffered saline (TBST) was performed on cryosections^23,24^.

Adult retinal flatmounts were performed by enucleating the right eye from fixed adult heads and processing the retinas for immunohistochemistry as previously described^25^. Primary antibodies included: goat α mCherry (AB0081-500, OriGene) 1:500, rabbit α Rbpms2 (AB181098-1001, Abcam) 1:1000, rabbit α Pax6 (12323-1-AP, Proteintech) 1:500, rabbit α Prox1 (AB5475, Millipore) 1:500, mIgG1 α Zpr1 (ANZPR-1, Zirc) 1:100, and wheat germ agglutinin (WGA) conjugated to Alexa 488 (W11261, ThermoFisher) 1:500. Secondary antibodies were all used at 1:500 and included: donkey α rabbit 488 (A21206, Invitrogen), donkey α mIgG1 488 (A21121, Invitrogen), and donkey α goat Cy3 (705-165-147, Jackson ImmunoResearch Laboratories, Inc). Nuclei were labeled with DAPI (4′,6-diamidino-2-phenylindole, 50 μg/mL) and slides cover slipped with Southern Biotech Fluoromount-G^®^.

For cleared tissue, we utilized the Accu-OptiClearing strategy, a protocol based on the OPTIClear clearing solution^26,27^. The reagents used were as previously described^27^. Adult wild type zebrafish were euthanized with cold 1X Tricaine and decapitated. The heads were fixed in 4% PFA/1X PBS overnight at 4°C. Heads were washed 3 times in 1X PBS on a rotator at room temperature, then whole brains, optic nerves and retina were dissected out as intact pieces, submerged in 4% SDS OPTIClear, and incubated at 37°C for two days until the tissue appeared mostly transparent. Samples were then washed 3 times with 1X PBS on a rotator at room temperature and protected from light. Brains were mounted in OPTIClear (no SDS) overnight at 37°C in the dark, using an 18 Chambered Coverglass System (Cellvis, C18-1.5H).

### Microscopy & Image Processing

5 dpf and adult cryosection images were taken on Zeiss AxioImagerZ.2, with the full adult retina sections stitched together using ZenBlue v. 3.5 software. Adult flatmount samples were imaged on a Nikon Laser scanning confocal. Cleared adult brain samples were imaged on the Andor BC43 and Dragonfly 620SR (with an inverted Leica DMi8 microscope) spinning disk confocal microscopes (Andor, Oxford Instruments) at the Oxford Instruments Center for Advanced Microscopy - Electron Microscopy Core (OxCAM-EM, RRID:SCR_026315) at the Medical College of Wisconsin. Cleared brain images were acquired with Fusion software (Oxford instruments) and processed using Imaris software (Oxford Instruments). Adult flatmount and brain images were processed in FiJi^28^. All images were processed using Adobe Photoshop. Cell counts were performed on one eye from each fish (n=5 or n=3) in the whole retinal section at all time points analyzed.

### Statistical Analyses

All data was analyzed on Graphpad Prism. Error bars on all figures represent mean ±standard deviation. Unpaired t-tests or one way ANOVA (analysis of variance) were run where appropriate (Prism) to determine statistical significance between experimental groups. *P* ≤ 0.05 was considered statistically significant.

## Results

### The atoh7:iCre transgene recapitulates endogenous atoh7 expression

To permanently label the zebrafish *atoh7* lineage, transgenic zebrafish expressing Cre recombinase from the zebrafish *atoh7* promoter were generated and crossed to the Cre inducible *ubi*:switch transgenic line^18^. In double transgenic offspring, cells containing Cre will switch their ubiquitous expression from eGFP to mCherry (Fig. 1A). To validate the *atoh7*:iCre expression was consistent, recapitulated endogenous expression, and did not produce leaky Cre expression, we characterized three separate founders.

**Figure 1.**
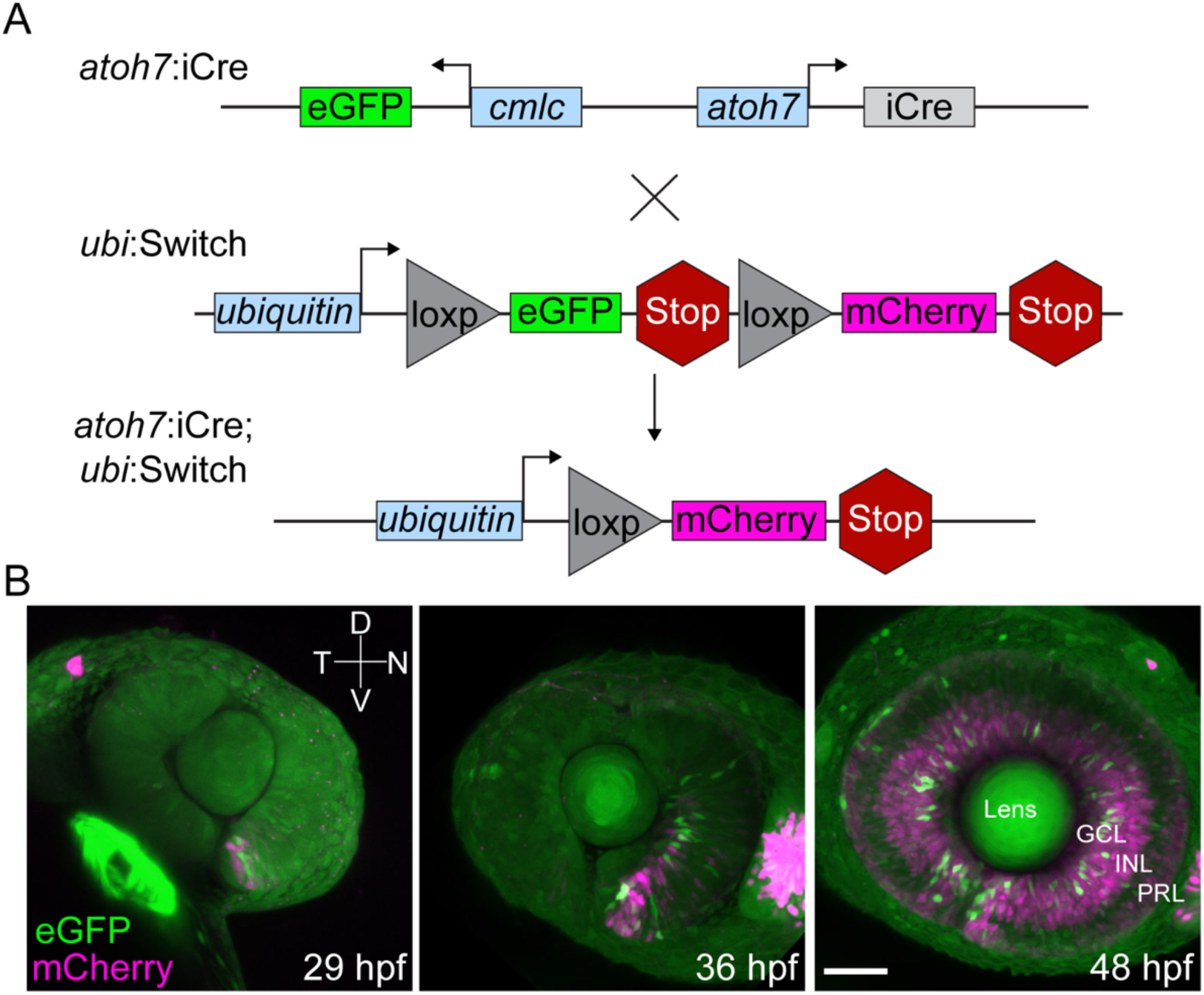
Timelapse *in vivo* imaging demonstrates permanent recombination of the *ubi*:Switch transgene. (A) Schematic of the *atoh7:*iCre;*cmlc:*GFP and *ubi:*loxp-eGFP-STOP-loxp-mCherry-STOP transgene cross resulting in embryos which are *ubi:*loxp-mCherry-STOP positive. (B) *In vivo* z-stack and maximum intensity projection of *atoh7:*iCre;*ubi:*Switch F2 larvae eyes at 29 hpf, 36 hpf, and 48 hpf respectively, demonstrating Cre recombination of *ubi*:Switch in a similar spatiotemporal pattern as endogenous *atoh7* expression. Scale bar 50μm. GCL, Ganglion Cell Layer; INL, Inner Nuclear Layer; PRL, Photoreceptor Layer.

Consistent recombination of the *ubi*:Switch transgene by all three founders was confirmed in 5 dpf larval retina sections (Fig. S2). Each founder demonstrated *atoh7*:iCre;*ubi*:Switch+ retinas contained mCherry+ cells in each retinal layer (Fig. S2). *ubi*:Switch or *atoh7*:iCre single positive retinas showed no mCherry or eGFP within the retina, respectively, confirming successful recombination and no leaky mCherry expression without Cre recombination (Fig. S1D).

Previous analysis of *atoh7* mRNA and transgene expression identified endogenous *atoh7* expression begins at 29 hpf in the ventral-nasal retina and spreads centrally/peripherally^12,14^. To confirm the expression of the *atoh7*:iCre transgene follows the same dynamic expression pattern as endogenous *atoh7,* we performed time-lapse confocal microscopy. As expected, mCherry expression became visible in the ventral-nasal retina by 29 hpf and spread centrally around the retina by 36 hpf (Fig. 1B). At 48 hpf, mCherry+ cells reached the temporal retina and spread into the periphery (Fig. 1B). Collectively, the *atoh7*:iCre transgene successfully recombined the *ubi*:Switch transgene in *atoh7+* cells within the developing zebrafish retina and showed a similar pattern to endogenous *atoh7* expression.

### Identification of atoh7 positive and negative RGCs in zebrafish

Lineage tracing analysis in mice found that ∼55% of mature RGCs come from the *atoh7*+ lineage despite the loss of ≥95% of RGCs in atoh7 mutants^5^. To investigate if zebrafish contain *atoh7* positive and negative populations of RGCs in wild type retinas, we quantified the colocalization of Rbpms2+ cells, which labels RGCs in the ganglion cell layer, and the *atoh7* lineage cells (mCherry+) in the F2 generation of all 3 *atoh7*:iCre founders^29,30^. At 5 dpf the *atoh7+* RGC population from each founder was 79% (± 3.60%), 82.4% (± 2.35%), and 86.6% (± 3.25%), while the *atoh7*-RGC averages are 21% (± 3.60%), 17.6% (± 2.35%), and 13.4% (± 3.25%), respectively (Fig. 2C, Fig. S2, Table S1). Due to similar averages in atoh7+ and - RGC lineage populations for each founder we chose to analyze one for the remainder of the experiments (founder 1). (Fig. 2C, Table S1).

**Figure 2.**
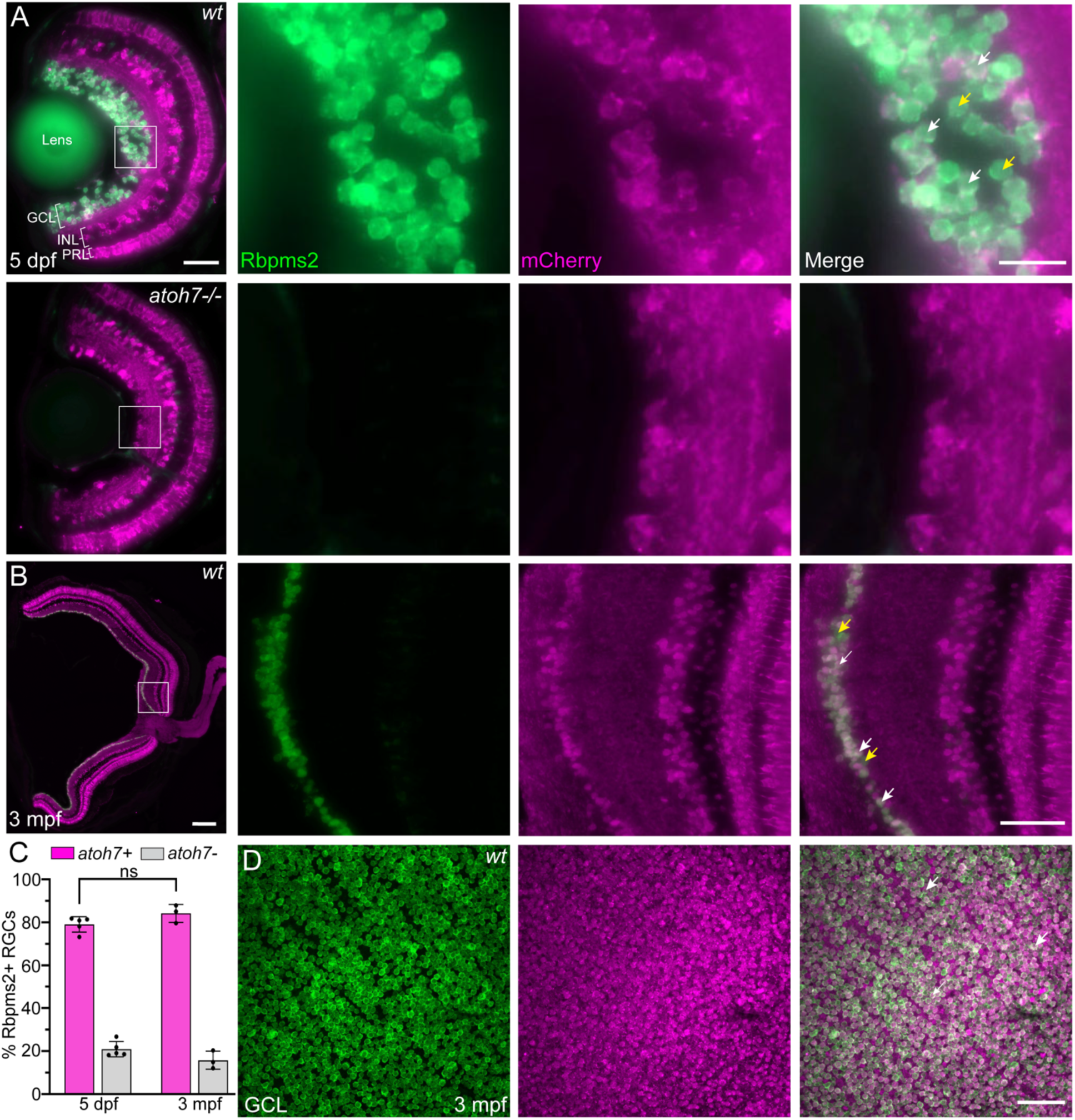
5 dpf and adult retinas show *atoh7* lineage positive and negative populations of RGCs. (A) Transverse sections (10μm) of 5 dpf *atoh7:*iCre;*ubi:*Switch larval retina with Rbpms2+ RGCs and *atoh7* lineage mCherry+ cells demonstrating the presence of *atoh7+* (white arrows) and *atoh7*-RGCs (yellow arrows). (B) Transverse sections (10μm) of 3-month (mpf) F2 *atoh7:*iCre;*ubi:*Switch retina with Rbpms2+ RGCs and mCherry+ *atoh7+* cells. Both *atoh7+* (white arrows) and *atoh7-* RGCs can be visualized. (C) Bar graph demonstrating no significant difference in the average percent of both *atoh7+* RGCs and *atoh7*-RGCs for 5 dpf (n=5) and 3 mpf (n=3) retinas (p=0.1154, two-tailed unpaired t-test). (D) Flatmount retina image from 3 mpf F2 *atoh7:*iCre;*ubi:*Switch retina focused on the GCL with Rbpms2+ RGCs and *atoh7* lineage mCherry+ cells. Scale bars, 50μm (A, left) and 20μm (A, right). ns, not significant, p ≥ 0.05. Error bars indicate standard deviation.

Since zebrafish eyes continue to grow throughout their life spans due to continued neurogenesis at the ciliary marginal zone (CMZ), we investigated if the proportion of *atoh7+* versus *atoh7-* RGCs remains consistent into adulthood. At 3 months post-fertilization (mpf), there was robust mCherry staining throughout all layers of the retina, similar to 5 dpf (Fig. 2B). In the GCL, 84% (± 4.2%) of RGCs were *atoh7+* and 16% (± 4.2%) *atoh7*-, consistent with the percentages observed at 5 dpf (Fig. 2C). In addition to the RGC population, there are Rbpms2-/mCherry+ cells in the GCL at 5 dpf and 3 mpf, which we speculate are displaced amacrine cells (Fig. 2A, B, D). Collectively, we show *atoh7* lineage positive and negative populations of RGCs are present as early as 5 dpf and their respective proportions remain consistent into adulthood.

### The zebrafish atoh7 lineage represents all major retinal cell types

The mouse *Atoh7* lineage includes all cell types derived from the multipotent RPC population, but it is unknown if this representation is conserved in the zebrafish *atoh7* lineage. In addition, *atoh7* mutant (*lakritz*) zebrafish retinas contain changes in the number or distribution of multiple cell types, including RGCs, amacrines, bipolars, and Müller glia, but it is unclear if these changes are reflected in the *atoh7* lineage population^6^. To determine the overall retinal population of *atoh7+* cells in wild type and *atoh7* mutants, we assessed the *atoh7+* and *atoh7-* populations using cell type specific markers.

As expected, *atoh7* mutants showed a complete loss of RGCs as indicated by an absence of Rbpms2 staining at 5 dpf (Fig. 2A), despite only 79% of RGCs being *atoh7+* in wild type retinas. (Fig. 2C and Table S1). However, we did see a consistent presence of mCherry+ cells within the GCL at 5 dpf, specifically in the dorsal region. To assess if these remaining mCherry+ cells are amacrine cells, we used Pax6, which labels amacrine cells and RGCs in the INL (inner nuclear layer) and GCL. These cells were Pax6+/Rbpms2-, suggesting they are displaced amacrine cells consistent with the previous analysis of *atoh7* mutant retinas (Fig. 3A, Fig. 2A)^6^. This data indicates both the *atoh7+* and *atoh7-* RGC populations are lost in *atoh7* mutants.

**Figure 3.**
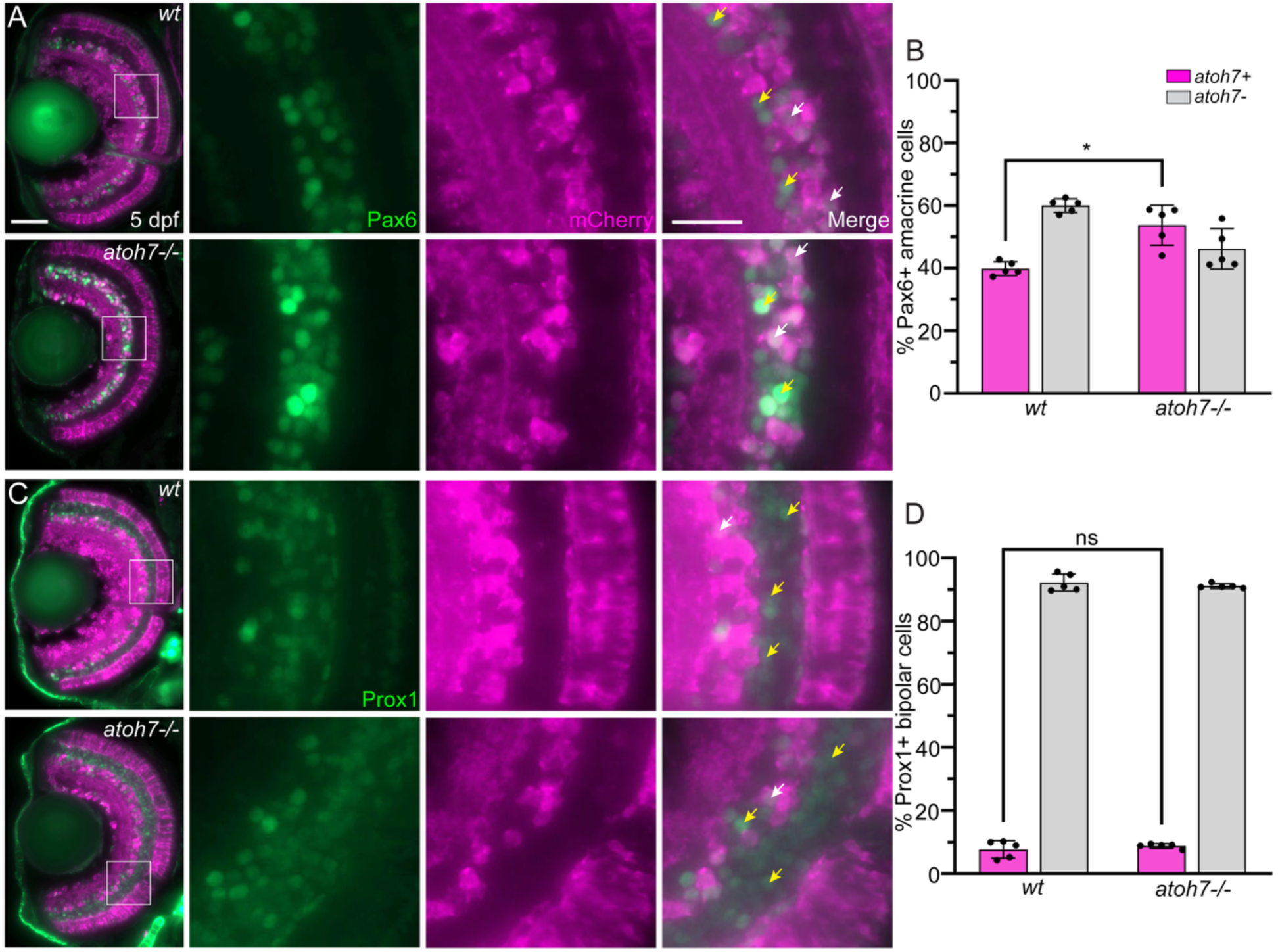
Amacrine cells, but not bipolar cells, have an increased *atoh7* lineage in *atoh7* mutants. (A) Transverse sections (10μm) of wild type (wt) and *atoh7-/-* 5 dpf *atoh7:*iCre;*ubi:*Switch larval retina with Pax6 antibody-stained amacrine cells in the INL and RGCs/amacrines in the GCL. White arrows indicate Pax6+ amacrine INL cells which colocalize with *atoh7* mCherry, while yellow arrows show Pax6+ amacrine INL that are *atoh7-*. (B) Bar graph demonstrating both *atoh7+* and *atoh7*-Pax6+ amacrine cell populations in wild type (n=5) and *atoh7 -/-* (n=5) backgrounds, showing a significant increase in the proportion of *atoh7+* cells in *atoh7 -/-* retinas compared to wild type (p=0.0018, two tailed unpaired t test). (C) Representative transverse section (10μm) images of 5 dpf *atoh7:*iCre;*ubi:*Switch wild type and *atoh7* mutant retinas stained with Prox1 to label bipolar cells. (D) Bar graph depicting percentage of *atoh7+* and *atoh7*-Prox1+ bipolar cell populations within the wild type (n=5) and *atoh7* mutant (n=5). Unpaired t-test analysis shows no significant difference in the *atoh7+* Prox1+ bipolar cell populations between genotypes (p=0.431, two tailed unpaired t test). Scale bars, 50μm (A) and 20μm (A), respectively. *, p < 0.05, ns, not significant, p ≥ 0.05. Error bars indicate standard deviation.

To determine the *atoh7+* amacrine cell lineage in the INL we counted Pax6+/mCherry+ and Pax6+/mCherry-cells in wild type and *atoh7* mutants. In wild type retinas, 40% (± 2.2%) of Pax6+ INL located amacrine cells come from the *atoh7* lineage whereas 60% (± 2.2%) are *atoh7*-(Fig. 3A & B, Table S1). Interestingly, in the *atoh7* mutants, 54% (± 6.4%) of Pax6+ INL amacrine cells are *atoh7+* and only 46% (± 6.4%) are *atoh7*-(Fig. 3A & B, Table S1), representing a significant increase in the number of *atoh7+*/Pax6+ cells in *atoh7* mutants compared to wild type. However, this increase in the proportion of *atoh7*+ amacrine cells was not reflected in the total amacrine cell counts in wild type (149 ± 27.62) and *atoh7* mutants (156 ± 29.38), indicating there was not an increase in the total INL amacrine cell population, but rather a shift from *atoh7-* to *atoh7*+ lineages (Table S1).

The *atoh7+* bipolar lineage in mice is <0.1% and slightly increases in *atoh7* mutants, accompanied by a decrease in the total bipolar cell population^5,31^. In the zebrafish retina the *atoh7+* bipolar population is 8%± 2.75%, marked by Prox1 and cellular location, a significantly greater proportion than mice (92% ± 2.75% were *atoh7*-). No change was observed in *atoh7* mutants (9% ± 0.76% *atoh7*+, 91% ± 0.76% *atoh7*-) despite the increase in the total bipolar cell population in *atoh7* mutants (288 ± 8.96) compared to wild type (134 ± 18.88) (Fig. 3C & D, Table S1). These results are consistent with previous zebrafish *atoh7* mutant characterizations, showing an increase in bipolar cells^6^. Collectively, this shows that in the absence of *atoh7*, there is not a shift in Prox1+ bipolar cells born from the *atoh7+* progenitor cell population and the wild type *atoh7*+ bipolar cell population in zebrafish is greater than in mice.

Müller glia (MG) cells are another small population represented in the mouse *atoh7+* lineage and differentially changed in *atoh7* mutant mice versus zebrafish^5–7,31^. In addition to RGCs, Rbpms2 is lowly expressed in MG cells in the zebrafish retina, evidenced by the expression discrepancy between RGCs and MG and morphology of Rbpms2 labeled cells in the INL^32–34^. Cell counts of Rbpms2+ MG found that the majority are *atoh7*-(99.5% ± 1.7%) compared to *atoh7*+ (0.5% ± 1.7%) and these numbers do not significantly change in *atoh7* mutants (99.4% ± 1.3% *atoh7*-, 0.6% ± 1.3% *atoh7*+) (Fig. 4A & B, Table S1). While previous reports described an increase in MG cells in *atoh7* mutant retinas we did not observe an increase in total MG cells in *atoh7* mutants (34 ± 2.28) compared to wild type (38 ± 7.3) (Table S1)^6^. Both mice and zebrafish MG cells are lowly represented in the *atoh7*+ population and the increase in MG observed in *atoh7* mutants are not from the *atoh7*+ lineage.

**Figure 4.**
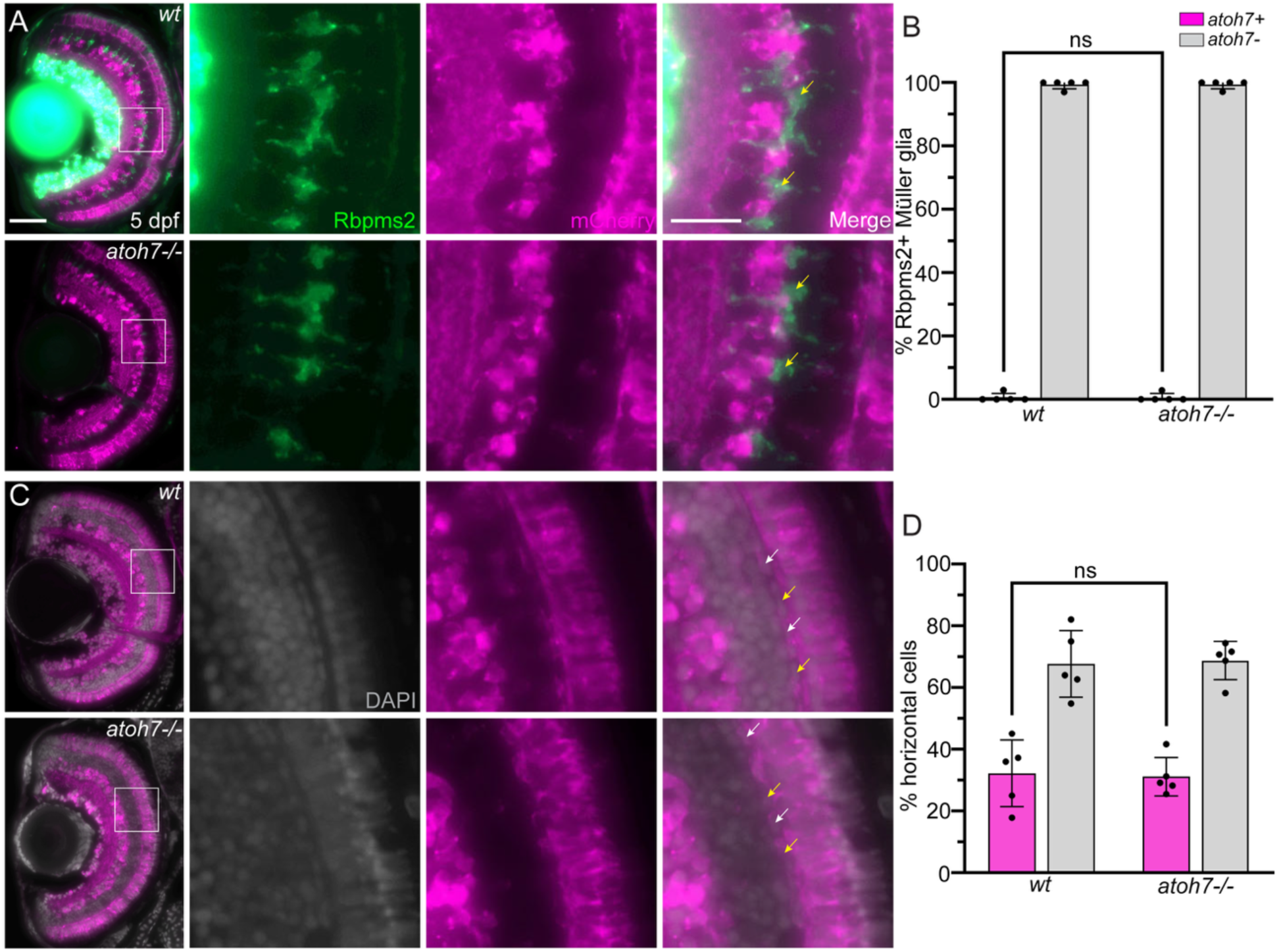
Müller glia and horizontal cells mostly arise from an *atoh7-* RPC lineage. (A) Transverse sections (10μm) of 5 dpf *atoh7:*iCre;*ubi:*Switch wild type and *atoh7* mutant retinas stained with Rbpms2 and overexposed to visualize Müller glia cells which are colocalized with *atoh7+* mCherry and those which are *atoh7*-(yellow arrows). (B) Percentage of *atoh7+,* Rbpms2+ Müller glia in both wild type (n=5) and *atoh7-/-*(n=5) show no significant difference in the proportion of *atoh7*+ Müller glia (p= 0.9842, two tailed unpaired t test). (C) 5 dpf *atoh7:*iCre;*ubi:*Switch wild type and *atoh7* mutant retinas with DAPI nuclear stain to highlight cell morphology. White arrows indicate *atoh7+* horizontal cells and yellow arrows indicate *atoh7*-cells. (D) Bar graph demonstrates *atoh7+* and *atoh7*-average percentages of horizontal cell lineages are not significantly different in wild type (n=5) compared to *atoh7* (n=5) retinas (p= 0.8526, two tailed unpaired t test). Scale bars, 50μm (A) and 20μm (A), respectfully. ns, not significant, p ≥ 0.05.

Next, we investigated the contribution of *atoh7+* progenitors to the horizontal cell population within 5 dpf retinas in both wild types and *atoh7* mutants. Utilizing DAPI to detect the distinct, oblong shape of horizontal cell nuclei at the apical portion of the INL, we identified 32% ± 10.8% of horizontal cells are born from the *atoh7+* lineage, while 68% ± 10.8% are *atoh7*-within wild type retinas (Fig. 4C & D, Table S1). Similar results were found in *atoh7* mutants; 31% ± 6.2% of horizontal cells are *atoh7+* and 69% ± 6.2% *atoh7*-(Fig. 4C & D, Table S1) with significant changes in the total number of horizontal cells in *atoh7* mutant (45 ± 5.1) compared to wild type retinas (53 ± 2.95) (Table S1). These results indicate there is no significant difference between *atoh7+* horizontal cell populations at 5 dpf in both wild type and *atoh7* mutants (Fig. 4C & D).

In the rod dominant mouse retina, loss of *Atoh7* results in increased cones and decreased rods, while in the cone rich zebrafish retina, cone numbers are reduced, which is similar to what occurs when *ATOH7* is reduced or lost in human retinas^7,31,35,35–38^. To determine the *atoh7*+ photoreceptor lineage in zebrafish and how it changes upon loss of *atoh7*, we used WGA to label rods and Zpr1 for red and green double cones. In wild type retinas, a majority or rods were *atoh7-* (77% ± 4.6%) compared to atoh7+ (23% ± 4.6 %) (Fig. 5A & B), while a large proportion of Zpr1+ cones came from the *atoh7*+ lineage (75% ± 3.2%) compared to *atoh7*-(25% ± 3.2%) (Fig. 5C &D). Within *atoh7* mutant retinas, neither the rod (21% ± 3.7%) or cone (76% ± 2.1%) *atoh7*+ lineage was significantly changed from wild type (Fig. 5, Table S1). Additionally, we did not observe any changes in the total number of rods or cones in the *atoh7* mutant retinas (90 ± 9.73 and 121± 4.12, respectively) compared to wild type (70 ± 5.76 and 112 ± 8.76, respectively) (Table S1). Overall, the zebrafish *atoh7+* photoreceptor lineages are higher than the reported percentages in mice.

**Figure 5.**
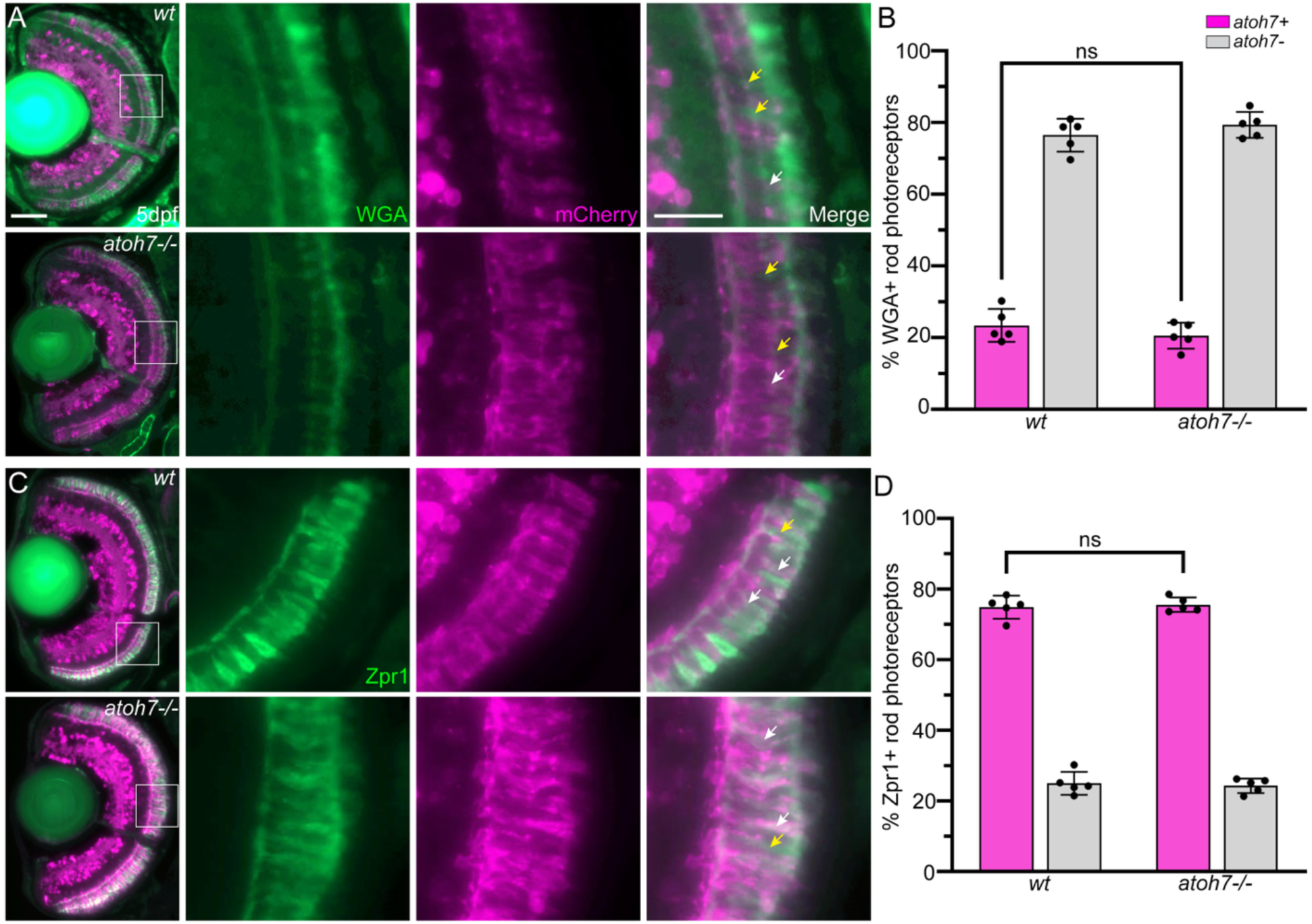
Most Zpr1+ cone photoreceptors are born from the *atoh7* lineage, but few WGA+ rod photoreceptors are *atoh7+*. (A) Transverse sections (10μm) of 5 dpf *atoh7:*iCre;*ubi:*Switch wild-type and *atoh7-/-* mutant retinas stained with WGA to identify the *atoh7+* (white arrows) and *atoh7-* (yellow arrows) rod photoreceptor population. (B) Bar graph representing percentage of *atoh7*+ (n=5) and *atoh7-* (n=5) WGA+ rod photoreceptors within wild-type and *atoh7-/-* retinas, demonstrating no significant shift in the proportion of the *atoh7* lineage in WGA+ rod photoreceptor cells (p=0.3077, two tailed unpaired t test). (C) Representative images of 5 dpf *atoh7:*iCre;*ubi:*Switch wild-type and *atoh7* mutant transverse retinal sections stained for *atoh7*+ cells (mCherry) and red-green double cone photoreceptors (Zpr1). *Atoh7*+/Zpr1+ cone photoreceptors are identified by white arrows and the *atoh7*-/Zpr1+ cones are identified with yellow arrows. (D) Bar graph representing no significant difference in the proportion of *atoh7*+ and *atoh7*-populations of Zpr1+ photoreceptors between wild-type (n=5) and *lakritz* (n=5) (p=0.6938, two tailed unpaired t test). Scale bars, 50μm (A) and 20μm (A), respectfully. ns, not significant, p ≥ 0.05.

### atoh7 lineage cells outside the zebrafish retina

*Atoh7+* cells and projections have been visualized outside of the retina^5,7,16,38–41^, so to thoroughly investigate the potential use of the *atoh7*:iCre transgene we analyzed cleared and immunostained adult brains at 3 months. As expected, we visualized cytoplasmic mCherry expression throughout the retina, optic nerve, optic chiasm, optic tract and nerve fibers superficially traveling across the optic tectum (Fig. 6A & C).

**Figure 6.**
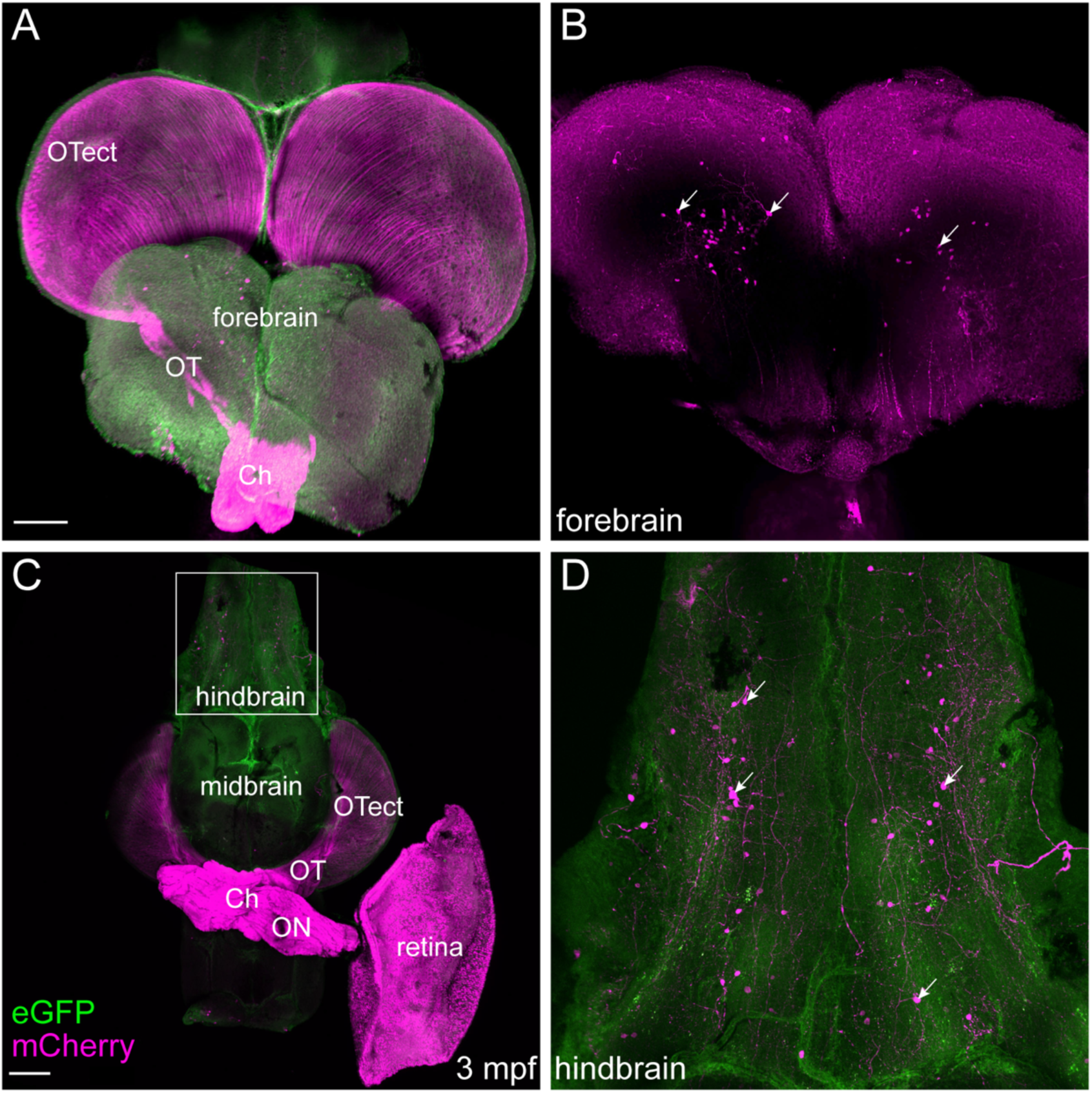
The *atoh7*:iCre transgene allows for identification of *atoh7+* cells in the central nervous system beyond the retina. (A) Frontal view of an *atoh7:*iCre;*ubi:*Switch adult (3 mpf) brain stained with a GFP antibody and mCherry antibody to enhance the signal of the *ubi:*Switch transgene and the *atoh7* lineage. (B) Transverse single plane maximum intensity projection within the middle forebrain region showing clusters of *atoh7+* cells bilaterally located within the deep, medial forebrain. (C) Ventral view of a whole mount *atoh7:*iCre;*ubi:*Switch cleared adult (3 mpf) brain stained with GFP and mCherry antibodies to enhance the signal of transgene expression. (D) Maximum intensity projection of hindbrain region showing populations of *atoh7+* cells present at the superficial, bilateral midline (white arrows). ON, Optic Nerve; Ch, Optic Chiasm; OT, Optic Tract; OTect, Optic Tectum. Scale bars, 300μm (A) and 200μm (C).

Outside the visual system we detected small populations of mCherry+ (*atoh7+*) cells within the deep, central forebrain (Fig. 6B) and observed *atoh7+* nuclei within the superficial ventral hindbrain (Fig. 6D). These regions of *atoh7+* cells are likely part of the auditory or olfactory processing centers, as *atoh7* has been shown to be expressed in the central auditory system of mice and olfactory placode in *Xenopus*^39–41^. Collectively, we demonstrate that our *atoh7*:iCre transgenic line alone or in combination with *ubi*:Switch can be used as a tool to investigate a variety of sensory system defects as they relate to the *atoh7* lineage or different mutations.

## Discussion

In combination with the *ubi*:Switch transgenic line, our *atoh7*:iCre transgenic zebrafish successfully labeled the *atoh7+* RPC population, allowing us to trace the lineage of transient *atoh7* expressing progenitors at time points when endogenous *atoh7* is no longer expressed. In the retina, we observed the presence of *atoh7+* and *atoh7-*populations of all major cell types. While presence of each retinal cell type within the *atoh7* lineage is consistent with the lineage tracing in mice, the *atoh7+* percentages differ except for horizontal cells and MG^5,31^. The differences observed between species may be due to the basic biology of each of these organisms. Mice are nocturnal and rely on navigating dimly lit environments and, thus, have a rod-dominant retina^42^. Zebrafish are diurnal and rely heavily on color vision to catch prey with a more cone-dominant retina^43^. The difference in photoreceptor cell composition is highlighted developmentally by their early production, with zebrafish cone photoreceptor precursors and RGCs being produced as daughter cells in the same mitotic division during the early stages of neurogenesis, a phenomenon not observed in mice^5,44^. This may explain why the proportion of *atoh7+* RGCs and cones were very similar (79 and 75%, respectively).

Outside of biology, technical differences such as Cre efficiency or transgene mosaicism, especially in a fast-vs slower-developing organism, may account for the *atoh7+* lineage differences between mice and zebrafish, although our ability to detect mCherry expression close to the onset of *atoh7* expression suggests developmental timing may not be an issue.

It was previously unknown if zebrafish contained an *atoh7-* RGC population due to the lack of a long-term lineage trace paradigm. Our interest in unveiling the zebrafish *atoh7* RGC lineage is due to the requirement of Atoh7 for the genesis and survival of a greater number of RGCs than is represented in the *Atoh7* lineage in mice, which creates two interesting questions: 1) how do *atoh7+* cells influence atoh7-RGCs and 2) what transcription factors and/or signaling pathways are responsible for *atoh7-* RGC genesis and survival. The data presented here reveals zebrafish contain an *atoh7-* RGC population providing support for the use of zebrafish to study the intricate RGC survival mechanisms important for nonautonomous RGC genesis and survival. In addition, our data demonstrates there was no change in the proportion of *atoh7+* RGCs in both the larval and adult retina, showing that as new cells are made at the CMZ, the *atoh7+* tRPCs generate RGCs in the same proportion as during development.

The *atoh7* lineage trace comparison between wild type and *atoh7* mutants validated previous results and revealed new information (Figure 7). In *atoh7* mutants the proportions of *atoh7+* cells for each cell type were not significantly changed, except in the Pax6+ amacrine cell population, which contains significantly more cells born from the *atoh7* lineage. Despite the increase in Pax6+/*atoh7+* amacrine cells, the total number of Pax6+ amacrine cells within the INL did not change, consistent with previous findings in *atoh7* mutants^6^. Our data also confirmed an increase in bipolar cells in *atoh7* mutants,^6^ although the increase was not associated with the *atoh7* lineage, suggesting the additional bipolar cells arise from an *atoh7*-progenitor population. Our MG cell count data in *atoh7* mutants was not consistent with previous data but the labeling method used differed(Table S1)^6^. Together, these results support the hypothesis that there is a fate switch of RPCs in *atoh7* mutants to later born cell types without the presence of Atoh7 protein to specify the RGC fate^5,6,31^.

**Figure 7.**
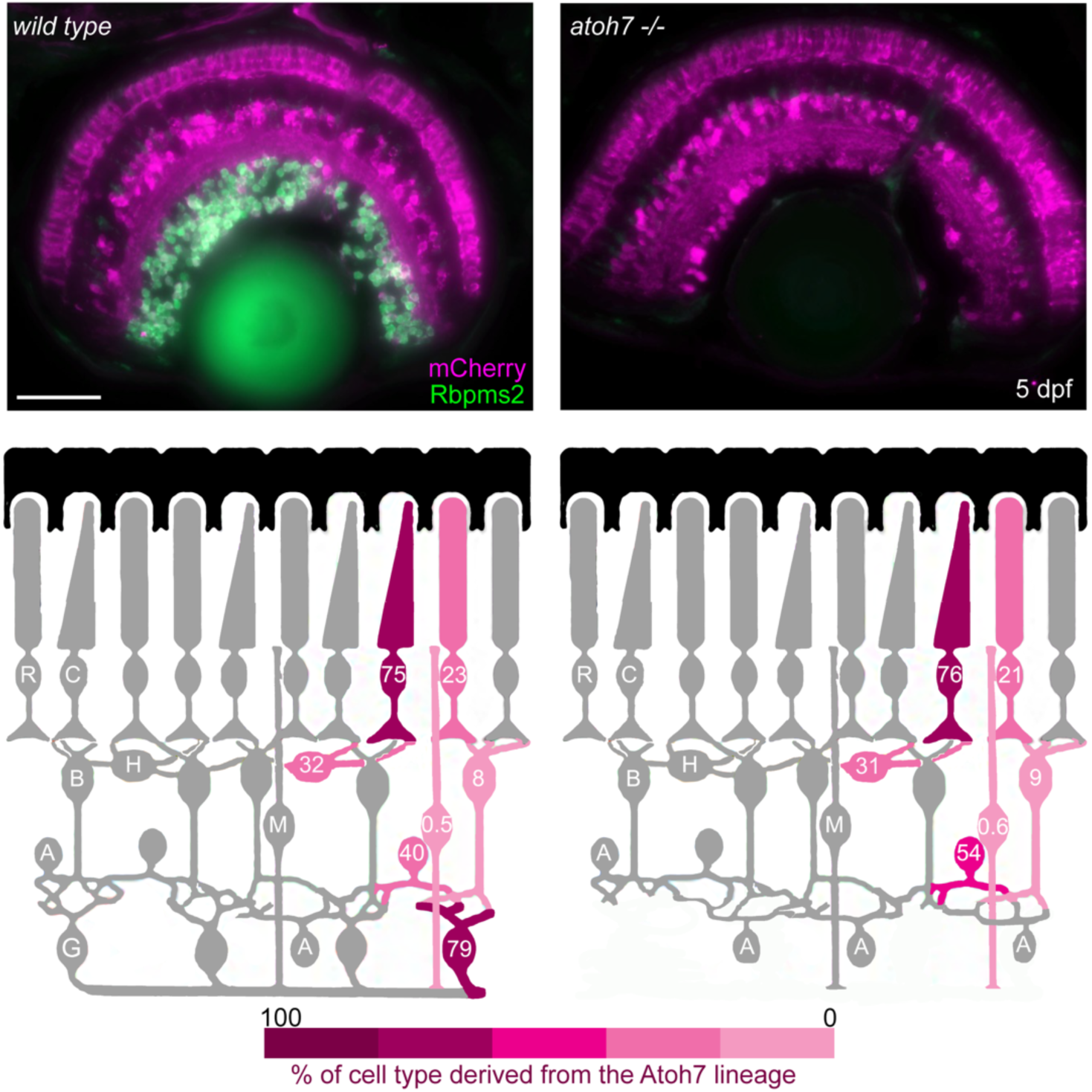
The *atoh7* lineage of each major cell type in the mature wild type zebrafish retina. A diagram depicting 5 dpf zebrafish retinal cell types and the respective proportions of each that are from the *atoh7* lineage in the retina of wild type and *atoh7* mutants. Quantified by colocalization quantifications of cell-specific stains and mCherry expression from the lineage tracing transgenes.

The influence of Atoh7 function and its lineage outside the eye in zebrafish has been limited due to the extra care needed to achieve adult survival of *atoh7* mutants. The lone publication investigating adult *atoh7* mutants reported a decrease in total brain area, although these results were agnostic to the specific regions of *atoh7* expression within the brain^35^. Providing support for the presence of *atoh7* activity outside of the retinofugal system, our cleared whole brain imaging of adult *atoh7*:iCre;*ubi*:Switch fish identified novel populations of *atoh7+* cells located within the deep, central forebrain and superficial ventral hindbrain, which may be related to deficits in total brain size as previously observed (Fig. 6A-D)^35^. The early expression of *atoh7* we observed in the developing olfactory bulb (Fig. 1B) and the expression in the adult brain compliments previous reports of *atoh7* being linked to the olfactory and auditory systems within *Xenopus* and mice^39–41^. Possible losses of olfactory neuron projections to areas in the central forebrain or auditory neuron projections to regions of the hindbrain may be what is potentially reducing brain size seen by others in *atoh7* mutants^35^.

Once championed as mainly a developmental model the use of zebrafish to study adult-onset human disease and regeneration of adult tissue has significantly increased over time. This change has prompted the need to extend our knowledge of zebrafish development, including the influence of the developmentally expressed transcription factor Atoh7. The generation of the *atoh7*:iCre lineage reporter, which recapitulates the timing of embryonic *atoh7* expression and can be detected into adulthood, confirmed previous reports and revealed new information. The existence of *atoh7* lineage positive and negative populations for each major retinal cell type (except for MG), including RGCs, provides opportunities to further explore RGC-specific survival mechanisms. The *atoh7*:iCre line is a unique tool to be utilized for lineage specific questions or targeted gene manipulations within the retina or CNS.

## Acknowledgements

We would like to thank Dr. Christian Mossimann for generously sharing the *ubi*:Switch transgenic line, Michael Cliff for zebrafish care, Dr. Brian Link for use of his confocal microscope and the Oxford Instruments Center for Advanced Microscopy - Electron Microscopy Core (RRID:SCR_026315) at the Medical College of Wisconsin, an institutionally available research service unit managed on behalf of the Medical College of Wisconsin by the Department of Cell Biology, Neurobiology and Anatomy.

## Commercial Relationships Disclosure

N/A

## Supplementary Material

**Supplemental Figure 1.**
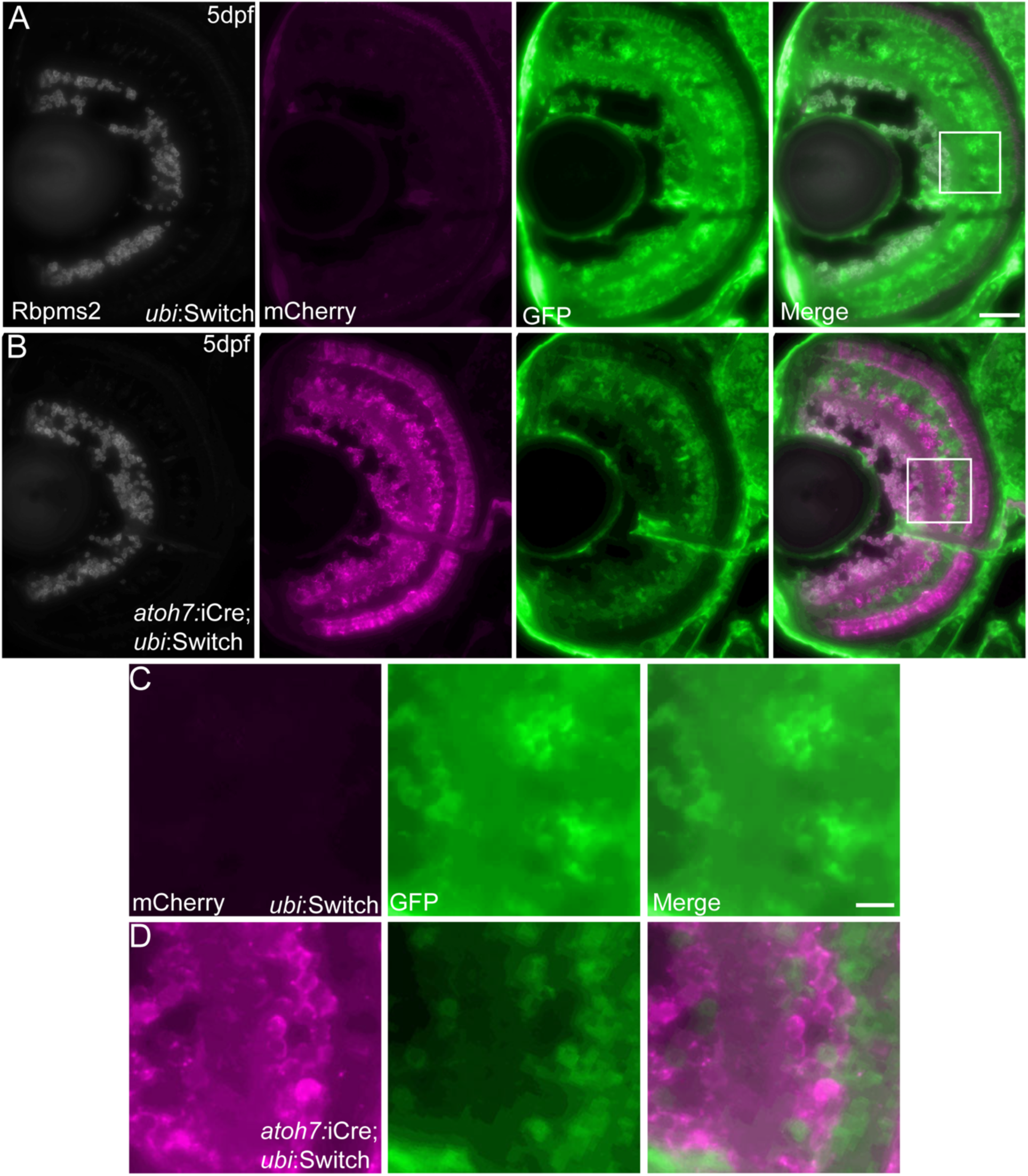
Successful quenching of transgene GFP fluorophore signal and anti-GFP antibody detection. (A) Transverse sections (10μm) of 5 dpf *ubi*:Switch larvae immunostained with Rbpms2 labeled RGCs and ubiquitous GFP (647 nm secondary detection) demonstrating that all cells of the retina are GFP+ without the Cre recombination. The GFP fluorophore is not detectable in the Rbpms2+ RGCs (488nm secondary detection) due to a MeOH series fluorophore quench. (B) Transverse sections (10μm) of 5 dpf *atoh7*:iCre;*ubi*:Switch larvae demonstrating quenching of GFP from the 488nm channel and detection of mCherry and Rbpms2. (C) Zoomed image of 5 dpf *ubi*:Switch retina demonstrating the absence of mCherry signal without Cre recombination and detection of GFP using an antibody in the 647nm channel. (D) Zoomed image of 5 dpf *atoh7*:iCre;*ubi*:Switch retina emphasizing the successful switching of the *ubi*:switch transgene without evidence of signal bleed through between 594nm and 647nm detection channels. Scale bars, 50μm (A, B) and 20μm (C, D), respectively.

**Supplemental Figure 2.**
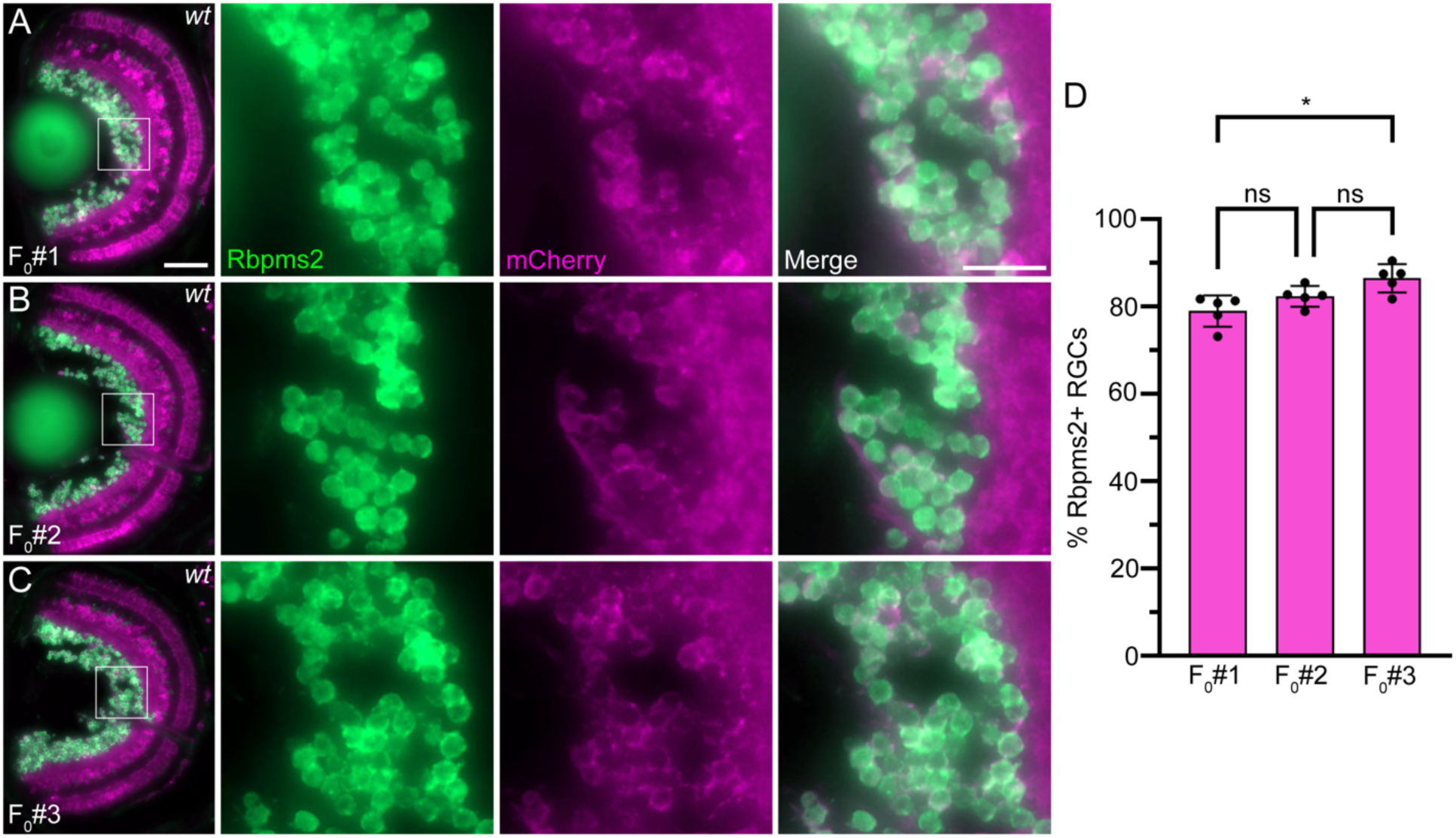
*Ubi*:Switch transgene showed specific recombination that was consistent in all founders. (A-C) Transverse sections (10μm) of 5 dpf *ubi*:Switch and *atoh7*:iCre;*ubi*:Switch F2 siblings from the founder 1 (A, F_0_#1), founder 2 (B, F_0_#2), and founder 3 (C, F_0_#3), highlighting Rbpms2 stainied RGCs (Rbpms2) and *atoh7+* cells (mCherry). Similar mCherry expression pattern was observed in all three founders. (D) Bar graph representing the percentage of *atoh7+* RGCs between all three of the founder lines by the second generation. This data shows that both F_0_#1 and F_0_#3 offspring are not significantly different from F_0_#2 offspring, but F_0_#3 offspring show significantly increased *atoh7+* RGCs compared to F_0_#1. Scale bars, 50μm (A, left) and 20μm (A, right), respectively. ns, not significant, p ≥ 0.05. *p ≤ 0.05 as determined with a one-way ANOVA with a Tukey’s multiple comparison test.

**Supplemental Table 1.** *Atoh7* lineage proportions for each cell type in both wild type and *atoh7* mutants. All cell types were either stained with cell specific antibodies or morphology and location were used in conjunction with mCherry colocalization with to determine the proportion of each cell type that came from the *atoh7* lineage. Each cell type count represents an average of n=5 retina from different larvae at 5 dpf (± standard deviation) for each cell type in both wild type and *atoh7* mutants.

| Cell Type | wild type |  |  | atoh7<sup>-/-</sup> |  |  |
| --- | --- | --- | --- | --- | --- | --- |
|  | atoh7<sup>+</sup><br>(a) | total<br>(b) | atoh7 lineage<br>(a/b x 100) | atoh7<sup>+</sup><br>(x) | total<br>(y) | atoh7 lineage<br>(x/y x 100) |
| RGCs | 178.40±14.79 | 225.80±18.54 | 79.01 | 0 | 0 | 0 |
| amacrine | 59.40±11.63 | 148.80±27.62 | 39.92 | 84.00±19.40 | 155.80±29.38 | 53.92 |
| bipolar | 10.40±3.85 | 134.40±18.88 | 7.74 | 25.60±2.51 | 288.40±8.96 | 8.88 |
| Müller glia | 0.20±0.45 | 37.60±7.27 | 0.53 | 0.20±0.45 | 33.8±2.28 | 0.59 |
| horizontal | 16.80±4.97 | 52.8±2.95 | 31.82 | 13.80±1.30 | 45.00±5.10 | 30.67 |
| rods | 16.20±2.28 | 69.8±5.76 | 23.21 | 18.60±4.34 | 89.80±9.73 | 20.71 |
| cones | 84.40±9.24 | 112.40±8.76 | 75.09 | 91.80±2.95 | 121.40±4.159 | 75.62 |

## References

1. Rapaport, D. H., Wong, L. L., Wood, E. D., Yasumura, D. & LaVail, M. M. Timing and topography of cell genesis in the rat retina. J Comp Neurol 474, 304–24 (2004).

2. Turner, D. L. & Cepko, C. L. A common progenitor for neurons and glia persists in rat retina late in development. Nature 328, 131–6 (1987).

3. Wong, L. L. & Rapaport, D. H. Defining retinal progenitor cell competence in Xenopus laevis by clonal analysis. Development 136, 1707–15 (2009).

4. La Vail, M. M., Rapaport, D. H. & Rakic, P. Cytogenesis in the monkey retina. J Comp Neurol 309, 86–114 (1991).

5. Brzezinski, J. A., Prasov, L. & Glaser, T. *Math5* defines the ganglion cell competence state in a subpopulation of retinal progenitor cells exiting the cell cycle. Dev. Biol. 365, 395–413 (2012).

6. Kay, J. N., Finger-Baier, K. C., Roeser, T., Staub, W. & Baier, H. Retinal Ganglion Cell Genesis Requires lakritz, a Zebrafish atonal Homolog. Neuron 30, 725–736 (2001).

7. Brown, N. L., Patel, S., Brzezinski, J. & Glaser, T. Math5 is required for retinal ganglion cell and optic nerve formation. Development 128, 2497–2508 (2001).

8. Wang, S. W. et al. Requirement for math5 in the development of retinal ganglion cells. Genes Dev. 15, 24–29 (2001).

9. Prasov, L. et al. ATOH7 mutations cause autosomal recessive persistent hyperplasia of the primary vitreous. Hum. Mol. Genet. 21, 3681–3694 (2012).

10. Ghiasvand, N. M. et al. Deletion of a remote enhancer near ATOH7 disrupts retinal neurogenesis, causing NCRNA disease. Nat. Neurosci. 14, 578–586 (2011).

11. Miesfeld, J. B., Glaser, T. & Brown, N. L. The dynamics of native Atoh7 protein expression during mouse retinal histogenesis, revealed with a new antibody. Gene Expr. Patterns 27, 114–121 (2018).

12. Kay, J. N., Link, B. A. & Baier, H. Staggered cell-intrinsic timing of*ath5*expression underlies the wave of ganglion cell neurogenesis in the zebrafish retina. Development 132, 2573–2585 (2005).

13. Brown, N. L., Dagenais, S. L., Chen, C.-M. & Glaser, T. Molecular characterization and mapping of ATOH7, a human atonal homolog with a predicted role in retinal ganglion cell development. Mamm. Genome 13, 95–101 (2002).

14. Masai, I., Stemple, D. L., Okamoto, H. & Wilson, S. W. Midline Signals Regulate Retinal Neurogenesis in Zebrafish. Neuron 27, 251–263 (2000).

15. Poggi, L., Vitorino, M., Masai, I. & Harris, W. A. Influences on neural lineage and mode of division in the zebrafish retina in vivo. J. Cell Biol. 171, 991–999 (2005).

16. Masai, I. et al. N-cadherin mediates retinal lamination, maintenance of forebrain compartments and patterning of retinal neurites. Development 130, 2479–2494 (2003).

17. Masai, I., Yamaguchi, M., Tonou-Fujimori, N., Komori, A. & Okamoto, H. The hedgehog-PKA pathway regulates two distinct steps of the differentiation of retinal ganglion cells: the cell-cycle exit of retinoblasts and their neuronal maturation. Development 132, 1539–1553 (2005).

18. Mosimann, C. et al. Ubiquitous transgene expression and Cre-based recombination driven by the *ubiquitin* promoter in zebrafish. Development 138, 169–177 (2011).

19. Zolessi, F. R., Poggi, L., Wilkinson, C. J., Chien, C.-B. & Harris, W. A. Polarization and orientation of retinal ganglion cells in vivo. Neural Develop. 1, 2 (2006).

20. Shimshek, D. R. et al. Codon-improved Cre recombinase (iCre) expression in the mouse. Genesis 32, 19–26 (2002).

21. Kwan, K. M. et al. The Tol2kit: A multisite gateway-based construction kit for Tol2 transposon transgenesis constructs. Dev. Dyn. 236, 3088–3099 (2007).

22. Kawakami, K. Transposon tools and methods in zebrafish. Dev. Dyn. 234, 244–254 (2005).

23. Brown, N. L. et al. Math5 encodes a murine basic helix-loop-helix transcription factor expressed during early stages of retinal neurogenesis. Development 125, 4821–4833 (1998).

24. Mastick, G. S. & Andrews, G. L. Pax6 Regulates the Identity of Embryonic Diencephalic Neurons. Mol. Cell. Neurosci. 17, 190–207 (2001).

25. Miesfeld, J. B. et al. The Atoh7 remote enhancer provides transcriptional robustness during retinal ganglion cell development. Proc. Natl. Acad. Sci. 117, 21690–21700 (2020).

26. Lai, H. M. et al. Next generation histology methods for three-dimensional imaging of fresh and archival human brain tissues. Nat. Commun. 9, 1066 (2018).

27. Lee, K. et al. Optimised tissue clearing minimises distortion and destruction during tissue delipidation. Neuropathol. Appl. Neurobiol. 47, 441–453 (2021).

28. Schindelin, J., et al. Fiji: an open-source platform for biological-image analysis. Nat. Methods 9, 676–682 (2012).

29. Hörnberg, H. et al. RNA-Binding Protein Hermes/RBPMS Inversely Affects Synapse Density and Axon Arbor Formation in Retinal Ganglion Cells In Vivo. J. Neurosci. 33, 10384–10395 (2013).

30. De Schutter, J. D. et al. Differential retinal ganglion cell resilience to optic nerve injury across vertebrate species. Front. Neurosci. 19, (2025).

31. Feng, L. et al. MATH5 controls the acquisition of multiple retinal cell fates. Mol. Brain 3, 36 (2010).

32. Farrell, J. A. et al. Single-cell reconstruction of developmental trajectories during zebrafish embryogenesis. Science 360, eaar3131 (2018).

33. Sur, A. et al. Single-cell analysis of shared signatures and transcriptional diversity during zebrafish development. Dev. Cell 58, 3028–3047.e12 (2023).

34. Sur, A. et al. Single-cell analysis of shared signatures and transcriptional diversity during zebrafish development. Dev. Cell 58, 3028–3047.e12 (2023).

35. Hammer, J. et al. Blind But Alive - Congenital Loss of atoh7 Disrupts the Visual System of Adult Zebrafish. Invest. Ophthalmol. Vis. Sci. 65, 42 (2024).

36. Lu, Y. et al. Single-Cell Analysis of Human Retina Identifies Evolutionarily Conserved and Species-Specific Mechanisms Controlling Development. Dev. Cell 53, 473–491.e9 (2020).

38. Brzezinski, J. A. th et al. Loss of circadian photoentrainment and abnormal retinal electrophysiology in Math5 mutant mice. Invest Ophthalmol Vis Sci 46, 2540–51 (2005).

39. Saul, S. M. et al. Math5 expression and function in the central auditory system. Mol. Cell. Neurosci. 37, 153–169 (2008).

40. Kanekar, S. et al. Xath5 participates in a network of bHLH genes in the developing Xenopus retina. Neuron 19, 981–994 (1997).

41. Burns, C. J. & Vetter, M. L. Xath5 regulates neurogenesis in the Xenopus olfactory placode. Dev. Dyn. 225, 536–543 (2002).

42. Joffe, B., Peichl, L., Hendrickson, A., Leonhardt, H. & Solovei, I. Diurnality and Nocturnality in Primates: An Analysis from the Rod Photoreceptor Nuclei Perspective. Evol. Biol. 41, 1–11 (2014).

43. Sato, S. & Kefalov, V. J. Characterization of zebrafish rod and cone photoresponses. Sci. Rep. 15, 13413 (2025).

44. Nerli, E. et al. Deterministic and probabilistic fate decisions co-exist in a single retinal lineage. EMBO J. 42, e112657 (2023).

